# Spatial Prior-Guided Boundary and Region-Aware 2D Lesion Segmentation in Neonatal Hypoxic Ischemic Encephalopathy

**DOI:** 10.1101/2025.03.13.642870

**Authors:** Amog Rao, Ananya Shukla, Jia Bhargava, Rina Bao, Yangming Ou

## Abstract

Segmenting acute and hyper-acute brain lesions in neonatal hypoxic-ischemic encephalopathy (HIE) from diffusion-weighted MRI (DWI) is critical for prognosis and treatment planning but remains challenging due to severe class imbalance and lesion variability. We propose a computationally efficient 2D segmentation framework leveraging ADC and ZADC maps as a three-channel input to UNet++ with an Inception-v4 encoder and scSE attention for enhanced spatial-channel recalibration. To address class critical imbalance and lack of volumetric context in 2D methods, we introduce a novel boundary-and-region-aware weighted loss integrating Tversky, Log-Hausdorff, and Focal losses. Our method surpasses state-of-the-art 2D approaches and achieves competitive performance against computationally intensive 3D architectures, securing a Dice Similarity Coefficient (DSC) of 0.6060, Mean Average Surface Distance (MASD) of 2.6484, and Normalized Surface Distance (NSD) of 0.7477. These results establish a new benchmark for neonatal HIE lesion segmentation, demonstrating superior detection of both acute and hyper-acute lesions while mitigating the challenge of loss collapse. The code is available at Neonatal-HIE-SPARSeg.

## 1 Introduction

Hypoxic ischemic encephalopathy (HIE) is a type of brain dysfunction (brain lesion injury) that occurs when the infant’s brain experiences a sudden decrease in oxygen or blood flow during the prenatal, intrapartum or postnatal period [10]. HIE affects around 1 to 5 per 1000 term-born infants, leading to significant long-term neurocognitive deficits such as developmental delays, cognitive impairment, cerebral palsy, epilepsy or death in about 60% of cases despite receiving therapeutic hypothermia treatment [3]. Accurate identification of brain lesions in neonatal MRIs is critical for prognosis, treatment evaluation, and understanding disease progression with the help of reliable biomarkers and detection methods. However, HIE lesions are often hyper-acute and multi-focal, posing challenges for algorithms that perform well on larger, focal lesions like brain tumours.

Apparent Diffusion Coefficient (ADC) maps help address this challenge by measuring the magnitude of water diffusion within a voxel of brain tissue and are calculated using Diffusion-Weighted MRI (DWI). The degree of reduction in ADC is correlated with the severity of the injury, indicating areas of decreased water diffusion. However, the normal range of ADC varies in space (different brain regions) and in time (as the brain develops rapidly during infancy), making expert interpretation error-prone [5,11]. For example, an ADC value of 800 (*×* 10^*−*6^ mm^2^*/*s) may be considered normal for cortical gray matter, where ADC values typically range from 780 *−* 1090 (*×* 10^*−*6^ mm^2^*/*s) [6]. However, the same ADC value in white matter, where normal ADC ranges from 620 *−* 790 (*×* 10^*−*6^ mm^2^*/*s), could be considered lesioned. This regional variability highlights the challenge of using absolute ADC thresholds for lesion detection.

Therefore, Weiss et al. proposed the Z-Score of ADC (Z_ADC_) to quantify deviations in a patient’s ADC value at each voxel relative to normative brain-specific range [19]. Normative ADC atlases, derived from healthy neonates, provide voxel-wise mean and standard deviation of ADC values. To enable direct comparison, the patient’s ADC map undergoes deformable registration, which aligns the patient’s brain scan with the atlas, ensuring an accurate anatomical correspondence across voxels. The Z_ADC_ value for each voxel *u* is calculated as,

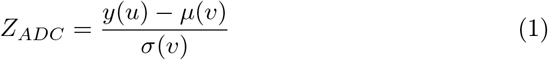

where, *y*(*u*) is the patient’s ADC value at voxel *u, µ*(*v*) is the mean ADC value at voxel *v* and *σ*(*v*) is the standard deviation of ADC at voxel *v*.

Demonstrating the utility of Z_ADC_ maps *∈* [*−*10, 10], comparisons can be made by thresholding them at values of *−*1.5, *−*2, and *−*2.5. Evaluating the accuracy of predicted masks using Dice Similarity Coefficient (DSC), sensitivity, and specificity, Bao et al. [3] demonstrated the highest DSC (0.54 *±* 0.28) at a threshold of *−*2. This threshold provided the best balance of accuracy and consistency in lesion detection across subjects, further highlighting the effectiveness of Z_ADC_ maps in lesion detection.

Given the current challenges of spatial variability and multi-focal nature of HIE lesions, Toubal et al. [4] integrated Swin-UNETR, a vision transformer-based segmentation model, with random forest classifier to enhance local feature discrimination. Their approach processes ADC and Z_ADC_ maps to generate a voxel-wise lesion probability map, segmented into overlapping 5 *×* 5 2D patches and classified using random forest. While effective for small datasets, the method is computationally intensive due to quadratic self-attention, 3D voxel-wise distance transforms, and sliding-window patch classification overhead.

Building on CNN architectures, Koirala et al. [8] proposed an ensemble of six 3D UNet variants, including dual-branch and attention-based architectures, with a hybrid loss combining BCE, MS-SSIM, and Jaccard Loss. Stratified 5-fold cross-validation, gradient accumulation, and sigmoid-averaged ensembling improved robustness and mitigated over-fitting. However, this approach incurs a high computational overhead arising due to multiple volumetric inferences and multi-scale loss computations, making training memory intensive.

In this work, we present a computationally efficient 2D segmentation framework for critically imbalanced, acute and hyper-acute lesion detection in neonatal HIE from sparsely sampled volumetric DWI. To achieve this we propose,

a. A three-channel input representation of ADC, Z_ADC_, and Z_ADC_ thresholded at *−*2 (Z_ADC_< *−*2) improving lesion detectability through enhanced diffusion-based feature representation.
b. A 2D framework, based on UNet++ with Inceptionv4-based encoder and scSE attention, leveraging both spatial and channel recalibration for improved feature extraction.
c. Tversky-Log-Hausdorff-Focal (TLHF) loss, a boundary and region-aware weighted loss designed to address loss collapse, where weak supervision suppresses gradient updates and leads to near-zero lesion predictions. TLHF improves spatial context, boundary delineation, and stabilizes gradient flow in highly imbalanced lesion segmentation.

Extensive ablation studies validate the effectiveness of our 2D framework, demonstrating competitive performance against computationally intensive 3D architectures, as evaluated on DSC, MASD, and NSD.

## 2 Dataset

The BOston Neonatal Brain Injury Dataset for Hypoxic-Ischemic Encephalopathy (BONBID-HIE) [2] is the first public, multi-center DWI dataset comprising of skull-stripped Apparent Diffusion Coefficient (ss-ADC) and Z-ADC maps with manually annotated lesion masks for 133 neonates. Patients had volumetric scan resolutions ranging from 128 *×* 128 *×D* to 256 *×* 256*×D*, where *D ∈* [16, 64]. Therefore, slice-thickness varies from (0.625mm *×* 0.625mm *×*2.0mm) to (2.0mm *×* 2.0mm *×* 6.0mm). The train set consisted of about 89 patients, whereas the test set consisted of 44 patients. Lesion volume analysis showed that 55.64% of patients had lesions occupying <1% of brain volume, 27.3% had 1%–5%, and 22.7% had >5%. The median lesion volume was 0.63% of total brain volume, indicating that DWI-detectable lesions in this cohort are typically hyper-acute.

## 3 Methodology

### 3.1 Pre-Processing

As shown in Figure 1, ADC, Z_ADC_, and a binary thresholded mask (Z_ADC_< *−*2) *∈ {* 0, 1*}* were stacked slice by slice along the axial axis. The thresholded mask was generated by applying a threshold of *−*2 to Z_ADC_, where pixels < *−*2 were given a positive label, while the remaining were assigned a negative label, creating a binary representation. These images were then resampled to a resolution of (3 *×* 256 *×* 256) using nearest-neighbors interpolation to prevent data distortion or addition of artifacts. To remove noise, ADC maps were clipped to eliminate negative pixel values captured during scanning. Next, min-max normalization was applied such that ADC [0, 3400] and Z_ADC_ [*−* 10, 10] were scaled to [0, 1], ensuring uniform pixel intensity for seamless channel stacking. To introduce diversity in input representations, non-aggressive augmentations such as random flips, gamma correction, anisotropic transformations and blurring were applied.

**Fig. 1.**
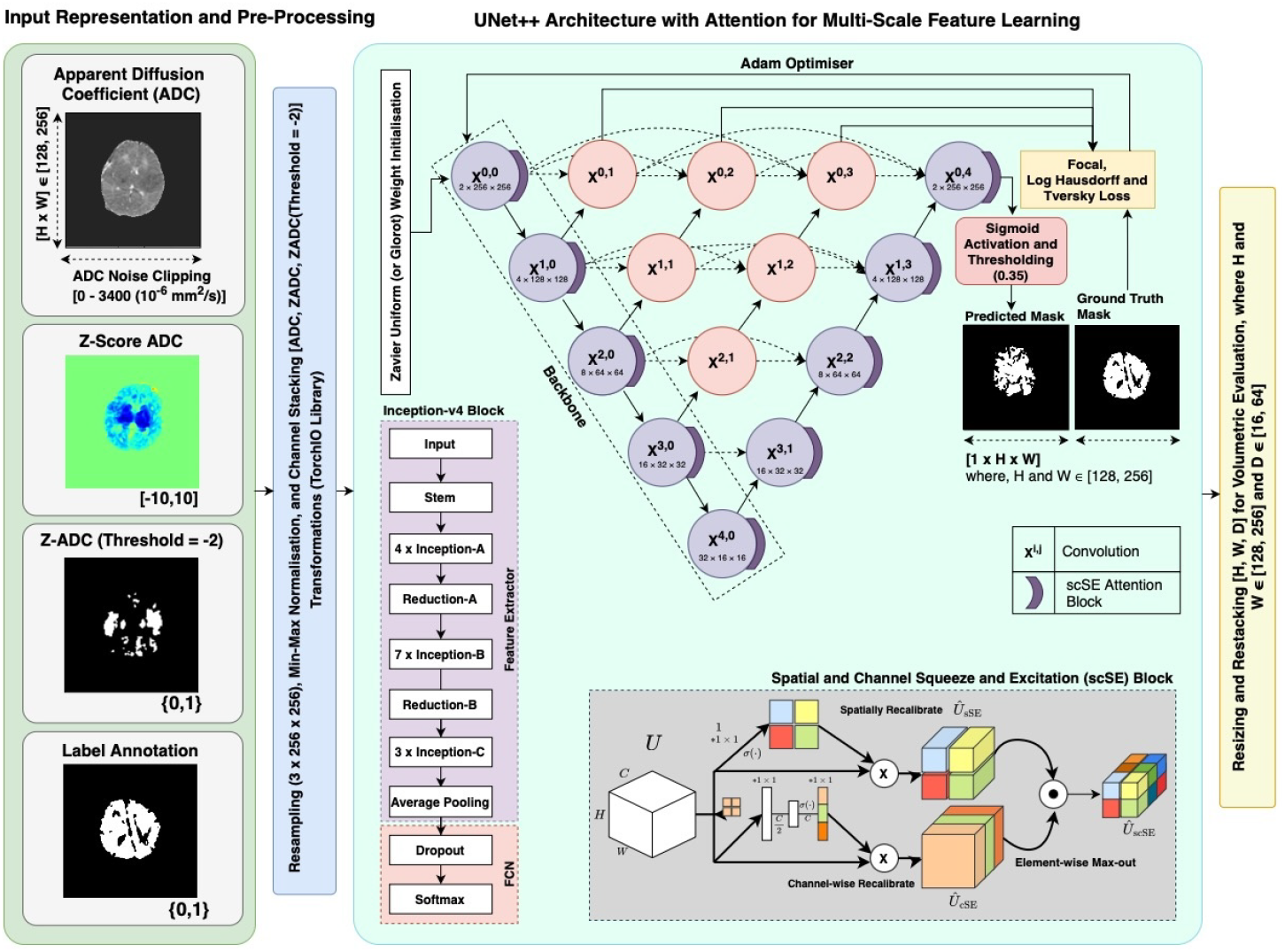
Methodological Pipeline.

### 3.2 Architecture

UNet++ [21] extends UNet [13] by introducing dense nested skip connections which progressively bridge the semantic gap between encoder and decoder feature maps. This enhances multi-scale feature aggregation and information flow. Additionally, deep supervision at multiple decoder stages ensures better gradient propagation, enabling the model to capture both global context and fine-grained features, which is critical for precise medical image segmentation. For 2D brain lesion segmentation, a three-channel input ADC - Z_ADC_ - Z_ADC_< *−*2 is stacked, normalized, and then fed into the Inception-v4 encoder blocks, which extracts multi-scale features through parallel convolutional paths, while 1×1 convolutions enable inter-channel mixing and spatial compression of feature maps [16].

Spatial and Channel Squeeze-and-Excitation (scSE) blocks [14] are applied after each encoder-decoder block to enhance feature representation. The Channel Squeeze-and-Excitation (cSE) branch applies global average pooling to compute channel-wise attention while, the Spatial Squeeze-and-Excitation (sSE) branch uses 1*×* 1 convolution and sigmoid activation to generate a spatial attention map, emphasizing pixel-level regions with higher importance. Concurrent recalibration by the cSE and sSE blocks, combined via element-wise addition, enhances lesion boundary delineation:

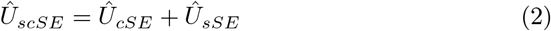

### 3.3 Loss Function

**Tversky Loss** [15] is a generalized Dice Loss (F-score equivalent) with tunable parameters *α* and *β* to control false positive and false negative penalties, improving performance on imbalanced datasets.

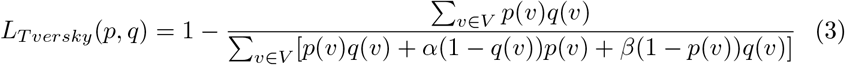

**Focal Loss** [9] is a variant of Cross-Entropy loss that focuses more on hard misclassified examples using a focusing parameter *γ*.

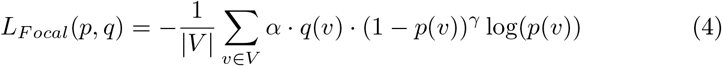

where, the weighting factor *α* balances the importance of positive and negative samples, while *γ ≥* 0 reduces the relative loss for easy-to-classify cases.

**Log-Hausdorff Loss** [7] captures boundary discrepancies by incorporating distance transforms (*d*_*p*_(*v*) and *d*_*q*_(*v*)), with a logarithmic term enhancing sensitivity to small errors.

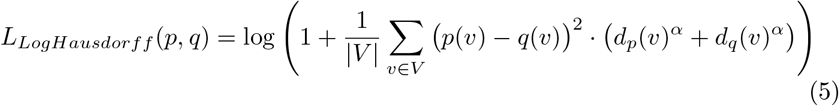

**Tversky-Log-Hausdorff-Focal Loss** - our customized boundary-and-region-aware loss, is a weighted combination of the above mentioned losses.

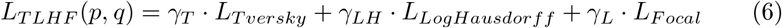

### 3.4 Hyperparameters and Training Strategy

Training was performed on an NVIDIA RTX A5000 (24GB) for 75–100 epochs with a batch size of 16. Model weights were initialized using Xavier uniform distribution, and optimization was handled via Adam with an initial learning rate of 0.0001, adjusted using ReduceLROnPlateau scheduler. TLHF Loss is a combination of Tversky (*α* = 0.3, *β* = 0.7, *γ*_*T*_ = 1.5), Log-Hausdorff (*γ*_*LH*_ = 2), and Focal (*γ* = 3, *γ*_*F*_ *∈ {* 2, 4 *}*) losses. Weight *γ*_*F*_ was dynamically adjusted across epochs to enhance sensitivity towards smaller lesions. Empirically, a threshold of 0.35 was identified as the optimal choice for converting the sigmoid outputs of the segmentation head into a binary mask.

## 4 Results and Inference

### 4.1 Evaluation Metrics

Segmentation accuracy is assessed using DSC [0, 1], MASD [0, *∞*), and NSD [0, 1] [12]. DSC measures global overlap, MASD quantifies distance from the region of interest, and NSD captures pixel-level misclassifications near boundaries. Together, these three metrics provide a robust measure of segmentation performance by ensuring overall region alignment and precise boundary localization.

**Dice Similarity Coefficient (DSC)** quantifies overlap between predicted (*p*) and ground truth (*q*) masks, given by

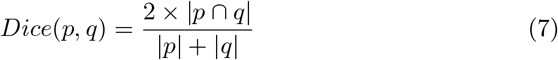

**Mean Average Surface Distance (MASD)** computes the average surface distance between the predicted mask (*p*) and ground truth (*q*), is given by

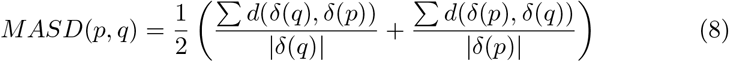

**Normalized Surface Distance (NSD)** measures surface overlap within a dilated tolerance *τ*,

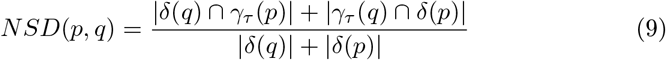

### 4.2 Results - Input Channel Stacking

This study, as indicated in Table 1 evaluated the impact of input channel stacking on segmentation performance using a UNet++ architecture with DenseNet- 161 and Inception-v4 encoders, trained with Dice-Focal loss. Models trained on a 3-channel input (ADC, Z_ADC_, and Z_ADC_< *−* 2) achieved the highest scores across all three metrics, significantly outperforming models trained on their 2- channel (ADC, Z_ADC_) counterparts, with DSC improvements of up to 5.98% for DenseNet-161 and 5.83% for Inception-v4. These results suggest that Z_ADC_< *−*2 masks act as spatial lesion priors, indicating regions with higher likelihood of lesion detection. Furthermore, Inception-v4 demonstrated superior performance across all metrics, due to its multi-scale feature extraction capability.

**Table 1.**
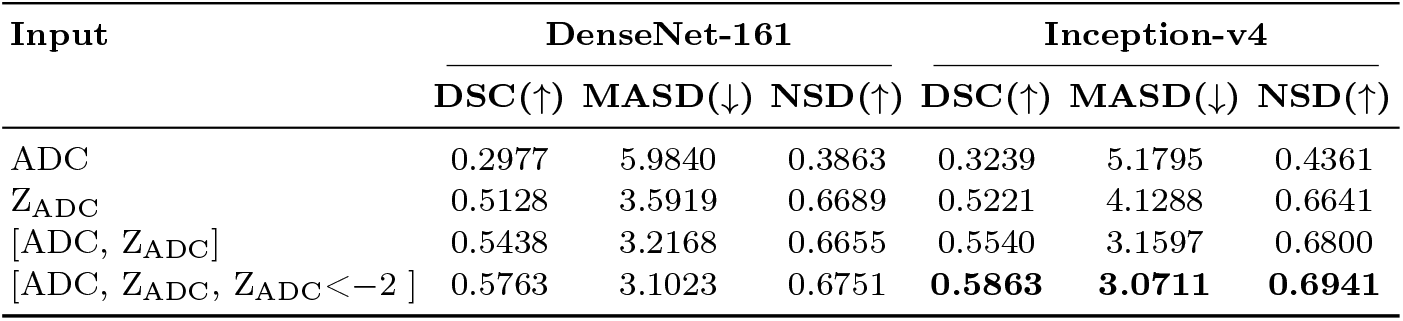
Ablation Study on Input Channel Stacking in DSC, MASD, NSD.

| Input | DenseNet-161 |  |  | Inception-v4 |  |  |
| --- | --- | --- | --- | --- | --- | --- |
| | DSC(↑) | MASD(↓) | NSD(↑) | DSC(↑) | MASD(↓) | NSD(↑) |
| ADC | 0.2977 | 5.9840 | 0.3863 | 0.3239 | 5.1795 | 0.4361 |
| $\text{Z}_{\text{ADC}}$ | 0.5128 | 3.5919 | 0.6689 | 0.5221 | 4.1288 | 0.6641 |
| [ADC, $\text{Z}_{\text{ADC}}$] | 0.5438 | 3.2168 | 0.6655 | 0.5540 | 3.1597 | 0.6800 |
| [ADC, $\text{Z}_{\text{ADC}}$, $\text{Z}_{\text{ADC}} < -2$] | 0.5763 | 3.1023 | 0.6751 | **0.5863** | **3.0711** | **0.6941** |

### 4.3 Results - Loss and Attention

As show in Figure 2 (a), Dice-Focal loss leads to loss-collapse, where lesion predictions shrink to near-zero, producing minimal gradients and trapping the model in a local minimum. Dice loss struggles with vanishing gradients, while Focal Loss, designed to address class imbalance amplifies the issue by suppressing false positives. Recovery occurs through stochastic perturbations, restoring gradients and stabilizing training. Large lesions (>5%) recover effectively due to stronger gradients, whereas smaller lesions (<1% and 1–5%) remain underrepresented, limiting effective supervision and refinement.

**Fig. 2.**
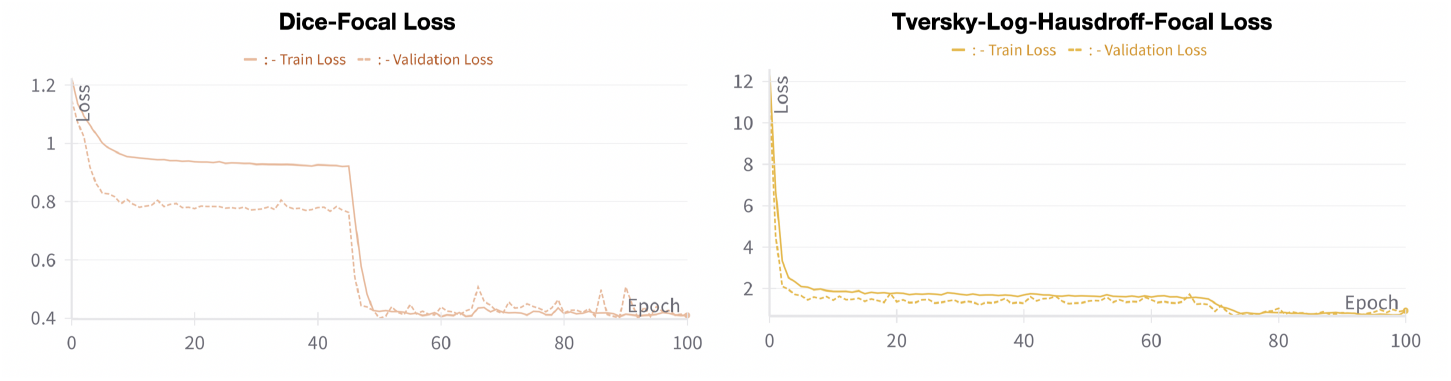
(a) Loss Collapse with Dice-Focal Loss (b) Mitigating Loss Collapse and using Tversky-Log-Hausdorff-Focal Loss.

To address loss collapse, we introduced a hybrid TLHF loss function combining boundary-based Log-Hausdorff loss, with region-specific Tversky loss, and class-balancing Focal loss. This approach enhanced training during early epochs, as indicated in Figure 2 (b), ensuring stable gradient flow and improved hyperacute lesion segmentation.

Performance of TLHF loss is evidenced in Tables 2 and 3, where it consistently outperforms Dice-Focal loss across all metrics. TLHF demonstrates improved lesion localization as indicated by MASD (2.6484), reduced boundary-based pixel-level misclassification through NSD (0.7477), resulting in a stronger global overlap captured by DSC (0.6060). Table 3 further confirms scSE’s impact, particularly for smaller lesions (<1%) where it significantly boosts NSD by 13%, making it more effective in capturing fine-grained structures.

**Table 2.** Ablation Study on Loss Functions and Attention Mechanisms.

| Loss | Attention | DSC (↑) | MASD (↓) | NSD (↑) |
| --- | --- | --- | --- | --- |
| Dice-Focal Loss | None | 0.5863 | 3.0711 | 0.6941 |
| TLHF Loss | None | **0.6060** | **2.6484** | 0.7153 |
| TLHF Loss | scSE | 0.5986 | 2.7986 | **0.7477** |

**Table 3.**
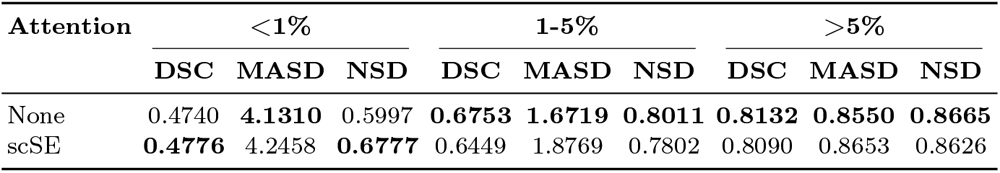
Best Performance Metrics Across Lesion Volume Percentages.

Our parameter-efficient 2D models, enhanced with scSE attention and trained on our boundary-and region-aware TLHF loss, achieve competitive performance in volumetric segmentation tasks despite the extensive parameters and contextual depth of 3D methods. Notably, our approach outperforms several 3D models, securing the second-highest NSD score and ranking in the top three for DSC, as shown in Table 4, demonstrating strong performance across existing studies on neonatal HIE lesion segmentation.

**Table 4.** Performance Evaluation across BONBID-HIE 2023 and 2024 Submissions.

| Participant/Team | Method | Architecture | DSC (↑) | MASD (↓) | NSD (↑) |
| --- | --- | --- | --- | --- | --- |
| Sejong_AI_team | 3D | UNet | 0.6262 | 2.5017 | 0.7316 |
| Toubal et al. [4] | 3D | Swin-UNETR w/ RF | 0.6215 | 2.2556 | 0.7678 |
| **Ours** | **2D** | **UNet++** | **0.6060** | **2.6484** | 0.7153 |
| **Ours** | **2D** | **UNet++ w/ scSE** | 0.5986 | 2.7986 | **0.7477** |
| Koirala et al. [8] | 3D | UNet Ensemble | 0.5800 | 2.5993 | 0.7379 |
| Wodzinski et al. [20] | 3D | ResUNet | 0.5770 | 2.9419 | 0.7379 |
| Tahmasebi et al. [17] | 3D | nnUNet | 0.4998 | 5.3327 | 0.6449 |
| Aydin et al. [1] | 3D | Attention UNet | 0.4826 | 3.4626 | 0.6176 |

## 5 Conclusion

In this work, we present a computationally efficient 2D segmentation framework for neonatal HIE lesions, leveraging a novel three-channel input including (ADC, Z_ADC_, and Z_ADC_ < *−* 2) as an anatomical prior to mitigate spatial and temporal uncertainties in DWI by encoding region-specific diffusion deviations. We introduce a hybrid region-and boundary-aware loss function, Tversky-Log-Hausdorff-Focal loss, that effectively addresses lack of spatial context in 2D methods and loss collapse in critically imbalanced datasets. Through our extensive ablation study, we demonstrate that spatial and channel attention mechanisms significantly enhance segmentation, particularly for hyper-acute (<1%) lesions. Our findings establish that our parametrically efficient 2D framework, outperforms resource-intensive 3D architectures, setting a new benchmark for neonatal HIE lesion segmentation. However, we acknowledge that 2D segmentation lacks volumetric context, and future work can explore ConvLSTM-based approaches [18] to model 3D data sequentially for improved spatial coherence.

## Acknowledgement

The authors acknowledge the invaluable guidance and computational support from Dr. Anupam Sobti, Plaksha University. This work was further enabled by the generous support of the organizers of the 2nd Boston Neonatal Brain Injury Dataset for Hypoxic-Ischemic Encephalopathy (BONBID-HIE) Lesion Segmentation and Outcome Prediction Challenge at MICCAI 2024, facilitating independent analysis.

## References

1 Aydin, M.A., Abdinli, E., Unal, G.: Segresnet based reciprocal transformation for bonbid-hie lesion segmentation. In: Bao, R., Grant, E., Kirkpatrick, A., Wachs, J., Ou, Y. (eds.) AI for Brain Lesion Detection and Trauma Video Action Recognition. pp. 39–44. Springer Nature Switzerland, Cham (2025)

2 Bao, R., Ou, Y.: BOston Neonatal Brain Injury Data for Hypoxic Ischemic Encephalopathy (BONBID-HIE): II. NICU and 2-Year Neurocognitive Outcome (2024). 10.5281/zenodo.13690270

3 Bao, R., Song, Y., Bates, S.V., Weiss, R.J., Foster, A.N., Jaimes, C., Sotardi, S., Zhang, Y., Hirschtick, R.L., Grant, P.E., Ou, Y.: BOston Neonatal Brain Injury Data for Hypoxic Ischemic Encephalopathy (BONBID-HIE): I. MRI and Lesion Labeling. Scientific Data 12(1), 53 (2025). 10.1038/s41597-024-03986-7

4 Eddine Toubal, I., Soltani Kazemi, E., Rahmon, G., Kucukpinar, T., Almansour, M., Ho, M.L., Palaniappan, K.: Fusion of deep and local features using random forests for neonatal hie segmentation. In: Bao, R., Grant, E., Kirkpatrick, A., Wachs, J., Ou, Y. (eds.) AI for Brain Lesion Detection and Trauma Video Action Recognition. pp. 3–13. Springer Nature Switzerland, Cham (2025)

5 Goergen, S., Ang, H., Wong, F., Carse, E., Charlton, M., Evans, R., Whiteley, G., Clark, J., Shipp, D., Jolley, D.: Early mri in term infants with perinatal hypoxic– ischaemic brain injury: interobserver agreement and mri predictors of outcome at 2 years. Clin Radiol 69, 72–81 (2014)

6 Helenius, J., Soinne, L., Perkiö, J., Salonen, O., Kangasmäki, A., Kaste, M., Carano, R.A., Aronen, H.J., Tatlisumak, T.: Diffusion-weighted mr imaging in normal human brains in various age groups. AJNR. American Journal of Neuroradiology 23(2), 194–199 (2002)

7 Karimi, D., Salcudean, S.: Reducing the hausdorff distance in medical image segmentation with convolutional neural networks. IEEE Transactions on Medical Imaging PP, 1–1 (07 2019). 10.1109/TMI.2019.2930068

8 Koirala, C.P., Mohapatra, S., Schlaug, G.: An ensemble approach for segmentation of neonatal hie lesions. In: Bao, R., Grant, E., Kirkpatrick, A., Wachs, J., Ou, Y. (eds.) AI for Brain Lesion Detection and Trauma Video Action Recognition. pp. 23–27. Springer Nature Switzerland, Cham (2025)

9 Lin, T.Y., Goyal, P., Girshick, R., He, K., Dollar, P.: Focal loss for dense object detection (10 2017). 10.1109/ICCV.2017.324

10 Long, M., Brandon, D.: Induced hypothermia for neonates with hypoxic-ischemic encephalopathy. Journal of Obstetrics Gynecology Neonatal Nursing 36, 293–298 (2007). 10.1111/j.1552-6909.2007.00150.x

11 Ozturk, A., Sasson, A., Farrell, J., Landman, B., da Motta, A., Aralasmak, A., Yousem, D.: Regional differences in diffusion tensor imaging measurements: assessment of intrarater and interrater variability. Am J Neuroradiol 29, 1124–1127 (2008)

12 Reinke, A., Eisenmann, M., Tizabi, M., Sudre, C., Radsch, T., Antonelli, M., Arbel, T., Bakas, S., Cardoso, M., Cheplygina, V., Farahani, K., Glocker, B., Heckmann-Notzel, D., Isensee, F., Jannin, P., Kahn, C., Kleesiek, J., Kurç, T., Kozubek, M., Landman, B., Litjens, G., Maier-Hein, K., Menze, B., Muller, H., Petersen, J., Reyes, M., Rieke, N., Stieltjes, B., Summers, R., Tsaftaris, S., van Ginneken, B., Kopp-Schneider, A., Jager, P., Maier-Hein, L.: Common limitations of image processing metrics: A picture story. ArXiv abs/2104.05642 (2021)

13 Ronneberger, O., Fischer, P., Brox, T.: U-net: Convolutional networks for biomedical image segmentation. In: Navab, N., Hornegger, J., Wells, W., Frangi, A. (eds.) Medical Image Computing and Computer-Assisted Intervention – MICCAI 2015, Lecture Notes in Computer Science, vol. 9351, pp. 234–241. Springer, Cham (2015). 10.1007/978-3-319-24574-4_28

14 Roy, A.G., Navab, N., Wachinger, C.: Recalibrating fully convolutional networks with spatial and channel “squeeze and excitation” blocks. IEEE Transactions on Medical Imaging 38(2), 540–549 (2019). 10.1109/TMI.2018.2867261

15 Salehi, S.S., Erdogmus, D., Gholipour, A.: Tversky loss function for image segmentation using 3d fully convolutional deep networks (09 2017). 10.1007/978-3-319-67389-9_44

16 Szegedy, C., Ioffe, S., Vanhoucke, V., Alemi, A.A.: Inception-v4, inception-resnet and the impact of residual connections on learning. In: Proceedings of the Thirty-First AAAI Conference on Artificial Intelligence. pp. 4278–4284. AAAI’17, AAAI Press (2017). 10.5555/3298023.3298188

17 Tahmasebi, N., Punithakumar, K.: A deep neural network approach for the lesion segmentation from neonatal brain magnetic resonance imaging. In: Bao, R., Grant, E., Kirkpatrick, A., Wachs, J., Ou, Y. (eds.) AI for Brain Lesion Detection and Trauma Video Action Recognition. pp. 34–38. Springer Nature Switzerland, Cham (2025)

18 Wang, Z., Sun, D., Zeng, X., Wu, R., Wang, Y.: Contextual embedding learning to enhance 2d networks for volumetric image segmentation. Expert Systems with Applications 253, 124279 (2024). 10.1016/j.eswa.2024.124279

19 Weiss, R., Bates, S., Song, Y., et al.: Mining multi-site clinical data to develop machine learning mri biomarkers: application to neonatal hypoxic ischemic encephalopathy. J Transl Med 17, 385 (2019). 10.1186/s12967-019-2119-5

20 Wodzinski, M., Müller, H.: Improving segmentation of hypoxic ischemic encephalopathy lesions by heavy data augmentation: Contribution to the bonbid challenge. In: Bao, R., Grant, E., Kirkpatrick, A., Wachs, J., Ou, Y. (eds.) AI for Brain Lesion Detection and Trauma Video Action Recognition. pp. 28–33. Springer Nature Switzerland, Cham (2025)

21 Zhou, Z., Siddiquee, M.M.R., Tajbakhsh, N., Liang, J.: Unet++: A nested u-net architecture for medical image segmentation. In: Stoyanov, D.e.a. (ed.) Deep Learning in Medical Image Analysis and Multimodal Learning for Clinical Decision Support, Lecture Notes in Computer Science, vol. 11045, pp. 3–11. Springer, Cham (2018). 10.1007/978-3-030-00889-5_1

